# Genomic normalization for sequencing libraries enrichment for rare somatic retroelement insertions

**DOI:** 10.1101/291773

**Authors:** Alexander Komkov, Gaiaz Nugmanov, Maria Salutina, Anastasia Minervina, Konstantin Khodosevich, Yuri Lebedev, Ilgar Mamedov

## Abstract

**Background:** There is increasing evidence that the transpositional activity of retroelements (REs) is not limited to germ line cells, but often occurs in tumor and normal somatic cells. Somatic transpositions were found in several human tissues and are especially typical for the brain. Several computational and experimental approaches for detection of somatic retroelement insertions was developed in the past few years. These approaches were successfully applied to detect somatic insertions in clonally expanded tumor cells. At the same time, identification of somatic insertions presented in small proportion of cells, such as neurons, remains a considerable challenge.

**Results:** In this study, we developed a normalization procedure for library enrichment by DNA sequences corresponding to rare somatic RE insertions. Two rounds of normalization increased the number of fragments adjacent to somatic REs in the sequenced sample by more than 26-fold, and the number of identified somatic REs was increased by 7.9-fold.

**Conclusions:** The developed technique can be used in combination with vast majority of modern RE identification approaches and can dramatically increase their capacity to detect rare somatic RE insertions in different types of cells.

## Background

In the past decade the rapidly growing number of whole genome sequencing studies proved the somatic variability to be the common property of genomes of both malignant and normal human cells (1-3). This somatic variability includes single nucleotide polymorphisms (SNPs), copy number variations (CNVs) and somatic insertions of active retroelements (REs) of L1, Alu and SVA subfamilies. Somatic RE insertions were found in several types of malignancies including lung, colorectal and prostate cancers (4-6). Studies of somatic RE insertions in normal cells were mainly focused on human brain since RE transpositions were shown to be associated with human adult neurogenesis (7-9). In other normal human tissues somatic RE variations are still poorly studied (10).

The modern experimental approaches for detection of somatic RE insertions is based on targeted high-throughput sequencing of genome fragments adjacent to RE insertions (TIP-Seq, RC-Seq, L1-Seq). However, even though the sequencing capacity of HTS technologies is growing rapidly somatic REs studies are still limited to few tissue samples, especially in case of low somatic insertions rate. At the moment, it is almost impossible to proceed the routine screening for somatic retroposition events in a sufficient number of individual cell genomes even using the most robust Illumina NovaSeq platform. Existing hybridization (14) and amplification-based enrichment techniques (13, 15) partially solve this problem allowing to increase the concentration of active RE subfamilies in sequencing libraries. Enrichment capacity achievable in these methods is sufficient to detect somatic RE insertions in most rapidly dividing cell samples such as tumor or embryonic cells where the proportion of somatic RE carrying cells is sufficient. However, somatic RE insertions (especially from large subgroups) presented in one or few cells of entire tissue sample remain almost undetectable among overwhelming majority of molecules corresponded to fixed and polymorphic ones. For instance, approximately 4,000 AluYa5 insertions are present in genomic DNA of each cell. Consequently, up to 800,000,000 molecules in AluYa5-enriched library represent fixed and polymorphic insertions in a 100,000 diploid cells sample whereas each somatic insertions can be presented in this sample by just several molecules. Thus, identification of rare somatic insertions without their specific enrichment is cost ineffective and looks like finding a needle in a haystack.

Another challenging point in somatic RE studies is the estimation of the number of cells in which a particular insertion is present. Most high-throughput sequencing library preparation techniques employ PCR amplification which inevitably introduce significant quantitative bias. As a result, the number of sequencing reads corresponding to each particular somatic insertion provides no assessment of the number of cells bearing this insertion even with usage of random fragmentation points for removing PCR duplicates.

Here we present the first approach for specific enrichment for rare somatic RE insertions in sequencing libraries. The method based on normalization procedure with utilization of Kamchatka Crab duplex-specific nuclease which allows to eliminate abundant DNA sequences and thus to increase the concentration of rare DNA sequences in the library. “Unique molecular identifiers” (UMIs) (16, 17) are used to remove PCR duplicates and estimate the true number of cells bearing a particular insertion. The method was employed for identification of AluYa5 somatic insertions in a sample of 50,000 nuclei from the adult human brain.

## Results

### The rationale of the method

The proposed method allows to identify rare somatic RE insertions (present in a single or few cells) using less sequencing reads. Furthermore, the method allows to quantify the number of cells that bear a particular insertion. There are three principal steps in the procedure:

i. Obtaining the genome fragments adjacent to RE insertions. In this study we performed selective amplification of the regions flanking retroelements of an evolutionary young AluYa5 subfamily using previously described technique (13, 18, 24, 25) with several modifications (see Fig 1 and selective amplification section below). Obtained amplicon contained sequences flanking AluYa5 insertion (about 90%) present in each cell, somatic AluYa5 insertion and sequences flanking insertions belonging to other Alu subgroups depleted during AluYa5-specific amplification. Sequences of non-Ya5 and somatic AluYa5 insertions were presented at a low level in the amplicon and were used for tracing changes of amplicon composition during subsequent normalization stages.
ii. Normalization using duplex-specific DNAse. At this stage, the amplicon is denatured and then slow renatured so that the abundant DNA molecules find their complementary pairs and return to the double-stranded (ds) state, while the rare molecules lag behind and remain single-stranded (ss). Subsequent treatment by duplex-specific DNAse from Kamchatka crab (26) eliminates dsDNA leaving ssDNA intact. After the amplification the relative abundance of molecules with low concentration in the original mix (including the flanks of somatic REs) is increased. This procedure is repeated twice to increase the enrichment efficiency.
iii. Sequencing of the normalized amplicons by Illumina and data analysis.

**Figure 1.**
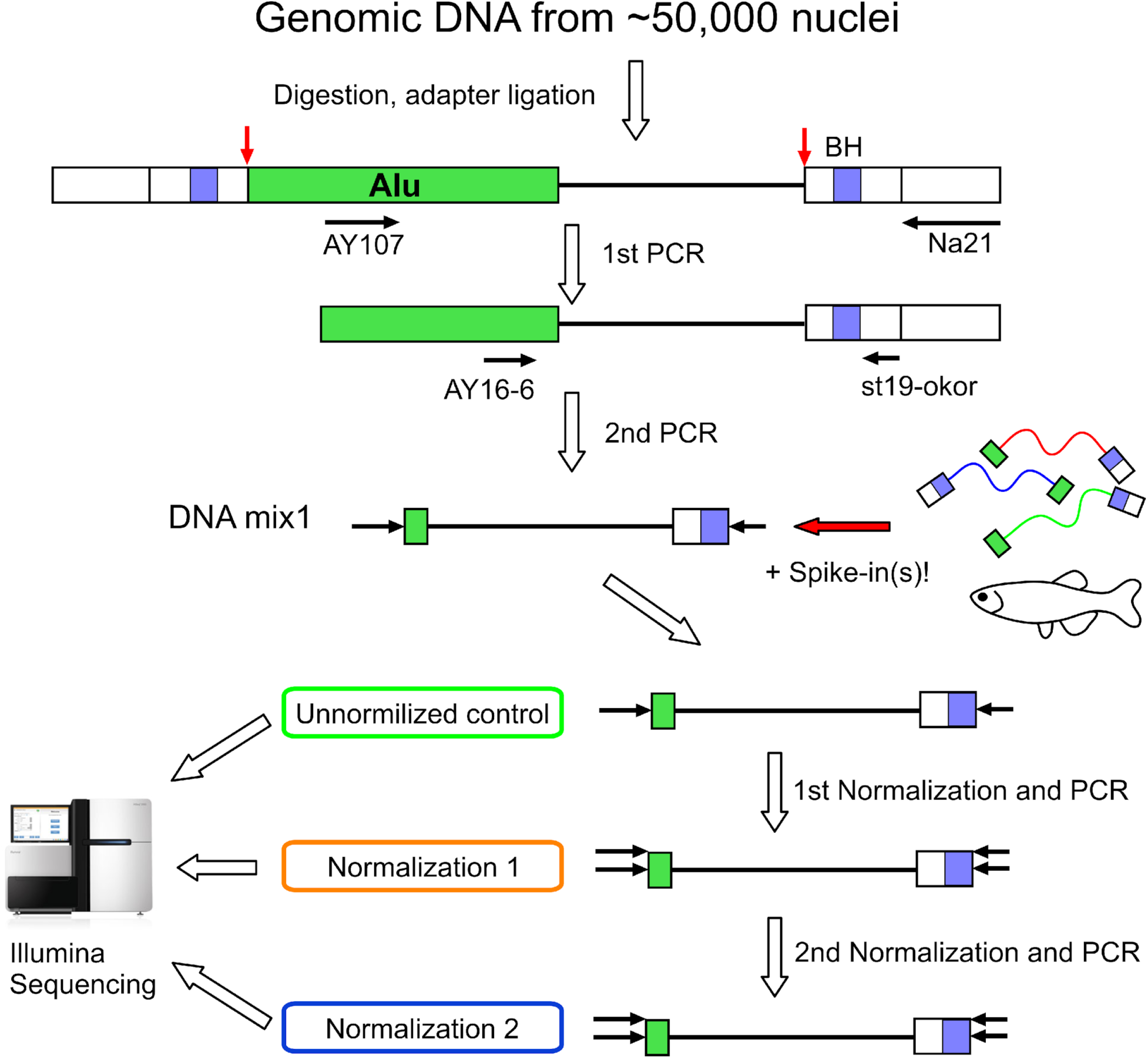
Overview of experimental procedure. Green boxes indicate Alu elements, white boxes – ligated adapter. Red arrows indicate genomic restriction sites for AluI, black horizontal arrows – primers and their annealing sites. Blue boxes (BH) – 8-nt molecular identifiers (MIs).

### Obtaining the genome fragments adjacent to RE insertions

50,000 nuclei were extracted from the frozen human brain sample (frontal cortex). Genomic DNA was extracted and used for selective amplification using suppression PCR. This procedure included DNA digestion by AluI endonuclease followed by ligation of suppressive adapters (see Fig. 1). Each molecule of the ligated adapter contains a “unique molecular identifier” (UMI) - a random sequence of 8 partly degenerated nucleotides (see Supplementary table 1 for oligonucleotide sequences). As a result, each of the ligated DNA molecules is marked by one of 6,561 different 8-nt oligomers prior to the amplification. UMIs allow to estimate the number of cells bearing a particular somatic insertion in case of sufficient sequencing depth. Sequences with identical UMI indicate a single ligation event and the number of different UMI correыponds to the number of cells containing each RE insertion. Following the adapter ligation two rounds of selective PCR were performed. In the first round, primer AY107 (25) was used for the selective amplification of insertions belonging to AluYa5 and AluYa8 subfamilies. The second primer anneals to the 5ʹ part of the ligated adapter. In the second round of amplification, a nested pair of primers was used: AY16-6 anneals to the 5ʹ end of an Alu element and St19okor primer to the middle part of the ligated adapter. As a result, each molecule in the amplicon contains two common parts at the ends (a 16 bp part of an Alu and a 27 bp adapter which includes the UMI) and a unique genomic flanking sequence for each insertion between (see Fig1) them.

**Table 1.**
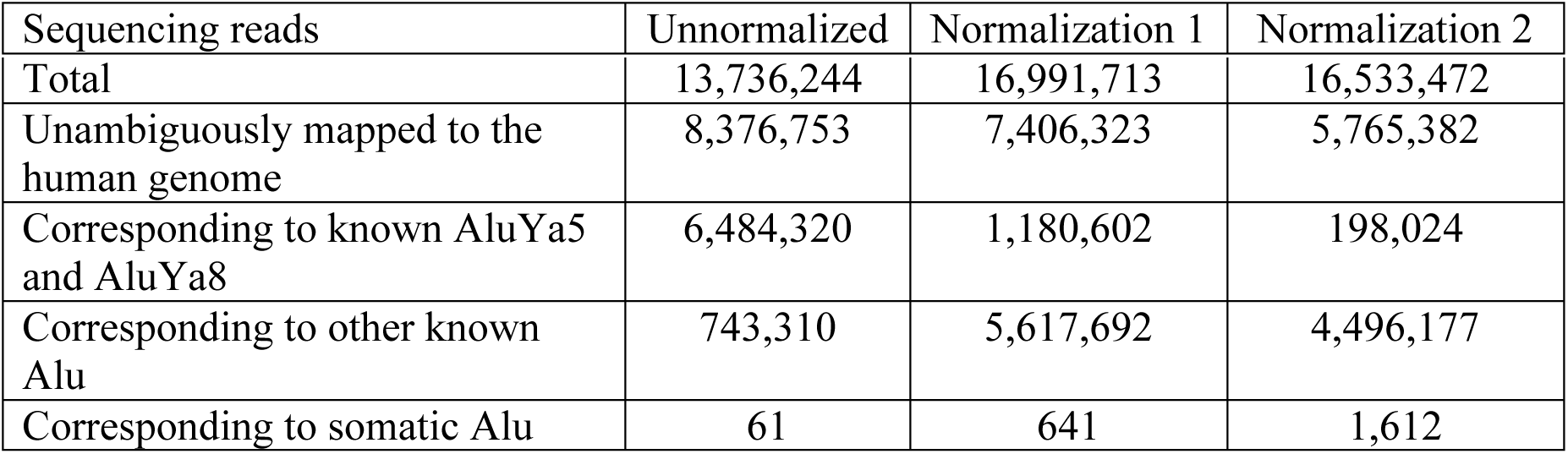
Distribution of sequencing reads

| Sequencing reads | Unnormalized | Normalization 1 | Normalization 2 |
| --- | --- | --- | --- |
| Total | 13,736,244 | 16,991,713 | 16,533,472 |
| Unambiguously mapped to the human genome | 8,376,753 | 7,406,323 | 5,765,382 |
| Corresponding to known AluYa5 and AluYa8 | 6,484,320 | 1,180,602 | 198,024 |
| Corresponding to other known Alu | 743,310 | 5,617,692 | 4,496,177 |
| Corresponding to somatic Alu | 61 | 641 | 1,612 |

### Spike-in controls

To monitor subsequent normalization, four artificial DNA fragments were added to the size-selected amplicon. These fragments ranging from 240 to 418 bp contain four different sequences from the genome of zebrafish (*Danio rerio*) which have the ends identical to those presented in all other fragments in the amplicon (a 16 bp part of an Alu and a 27 bp adapter introduced by step-out PCR). Two of these fragments (240 bp and 389 bp in length) were added in a concentration corresponding to a somatic insertion that is presented in five out of 50,000 cells whereas two other (259 bp and 418 bp in length) in the concentration corresponding to an insertion that is presented in one out of 50,000 cells (see Materials and Methods). Following the addition of spike-in controls, the mixture was divided into two equal aliquots. One aliquot was sequenced and used as unnormalized control whereas the other one was subjected to normalization using duplex-specific endonuclease.

### Normalization using the duplex-specific endonuclease

The amplicon was denatured, renatured and treated by the duplex-specific endonuclease. During renaturation DNA fragments with high concentration find their complementary chains and anneal to form dsDNA whereas fragments with low concentration remain single-stranded in the mix. As a result of subsequent digestion by duplex-specific DNAse, the majority of highly abundant fragments (corresponding to fixed AluYa5 insertions) were digested whereas rare fragments (including somatic AluYa5 insertions, spike-in controls and previously depleted other Alus such as AluYb8) remained intact. The normalized amplicon was reamplified with the primers used for the second round of selective amplification (AY16-6/St19okor) and again split to two equal portions. The first portion (“normalization 1”) was ligated to the Illumina adapters and sequenced. The second portion was subjected to second round of normalization, reamplified (“normalization 2”), ligated to the Illumina adapters and sequenced.

### Sequencing and data analysis

Three libraries (“unnormalized”, “normalization 1” and “normalization 2”) were sequenced using Illumina HiSeq. More than 47 millions of sequencing reads were obtained (see Table 1 for details). The vast majority of reads in the “unnormalized” library represented the sequences flanking AluYa5 insertions. About 80% of reads represented known AluYa5 insertions (annotated in Human Genome Browser, in databases of polymorphic REs), while 11% of sequences corresponded to the flanks of polymorphic or germline AluYa5 insertions found in the genome of the same donor in our previous study (13). About 9% of sequencing reads originated from the Alu insertions of other subfamilies. The Alu subfamily composition of normalized libraries significantly changed as a result of the normalization process (Table 1). The identification of potentially somatic insertions was performed as previously described (13, 18). Briefly, all sequencing reads were mapped to the reference human genome and the obtained coordinates were compared to the coordinates of fixed and polymorphic Alu insertions. To filter out the insertions present in all tissues of the donor, the remaining coordinates were compared to the previously identified Alu coordinates from four other tissues of the same individual (13). Only the insertions that did not match any RE insertion in the human genome and were absent from the other four tissues of the same individual were considered potentially somatic. Additionally all artificial sequences (e.g. chimeric reads, PCR fragments resulting from mispriming, etc) were filtered out using previously described stringent algorithms (18). Genomic coordinates, sequencing reads and the distribution of UMIs is shown in Supplementary table 2.

**Table 2.** Number of sequencing reads and UMIs corresponding to somatic insertions and spike-in controls

|  | Unnormalized | Normalization 1 | Normalization 2 |
| --- | --- | --- | --- |
| Sequencing reads corresponding to somatic Alu insertions | 61(51) * | 641(227) * | 1612(468) * |
| Number of potential somatic insertions | 48 | 173 | 379 |
| Number of potential somatic insertions with UMI count > 1 | 1 | 26 | 39 |
| Spike-in controls |  |  |  |
| DR240 (in 5 cells**) | 0 | 4(3)* | 7(4)* |
| DR389 (in 5 cells**) | 1(1)* | 1(1)* | 1(1)* |
| DR259 (in 1 cell***) | 0 | 0 | 1(1)* |
| DR418 (in 1 cell***) | 0 | 0 | 0 |

\* number of MIs is given in parentheses
\*\* corresponds to an insertion present in 5 out of 50,000 cells
\*\*\* corresponds to an insertion present in 1 out of 50,000 cells

### Evaluation of the method efficiency for library enrichment for somatic RE insertions

The efficiency of normalization was evaluated by direct counting of the number of somatic insertions, sequencing reads and UMIs corresponding to somatic insertions and spike-in controls (see Table 2). The number of identified somatic insertions increased more than 3.5-fold (from 48 to 173) after the first round of normalization and 7.9-fold (from 48 to 379) after the second round compared to the “unnormalized” library. Pearson’s Chi-squared test indicated a significant increase in the proportion of somatic insertions relative to fixed ones (*p* = 9.7×10^−5^ for “unnormalized” versus “normalization 1”; *p* = 4.5×10^−13^ for “normalization 1” versus “normalization 2”; *p* < 2.2×10^−16^ for “unnormalized” versus “normalization 2”). The number of sequencing reads representing somatic insertions increased from 61 in “unnormalized” library to 641 and 1,612 after the first and the second rounds of normalization respectively. 39 out of 379 insertions identified in the “normalization 2” library had more than one UMI indicating that these insertions were initially present in more than one cell. Only one out of four spike-in controls was detected in the “unnormalized” library. Two spike-in controls were identified in the “normalization 1” library whereas three out of four spike-in controls were detected in the “normalization 2” (see Table 2). The number of sequencing reads corresponding to spike-in controls also increased from one in the “unnormalized” to nine in the “normalization 2” library.

We additionally employed quantitative PCR (qPCR) as another method to estimate efficiency of normalization. To this end, we used primer pairs that corresponded to sequences flanking four fixed AluYa5 insertions, four randomly selected somatic insertions having more than one UMI and four spike-in controls (Fig2 and Supplementary table 3). The qPCR data indicated that the concentration of fixed AluYa5 insertions decreased by approximately 4-30 fold after the first round of normalization and by 8-30 fold after the second round (Fig 2, red dots). Oppositely, the concentration of spike-in controls increased by 8-30 times for the ones added in concentration of five cells and by 130-250 times for the sequences added at concentration corresponding to one cell per 50,000. Thus, the increase in the concentration of spike-in controls depended on the initial abundance in the amplicon before normalization. After the second round of normalization, the concentration of spike-in controls additionally increased by 2-8 times. (Fig 2, green dots). Furthermore, the selected somatic insertions initially presented at higher concentrations compared to the spike-in controls were also significantly enriched in the course of normalization (Fig 2 blue dots). Thus, the ratio between highly abundant and rare sequences of the initial amplicon was greatly decreased by normalization leading to more universal distribution of RE frequencies in the amplicon. Strikingly, as shown in Fig 2, the difference between the most abundant and the most rare sequence in our experiment changed from nearly 25 qPCR cycles (that is roughly 33,000,000-fold difference in concentration) to only 10 cycles (corresponding to 1,000-fold concentration difference).

**Figure 2.**
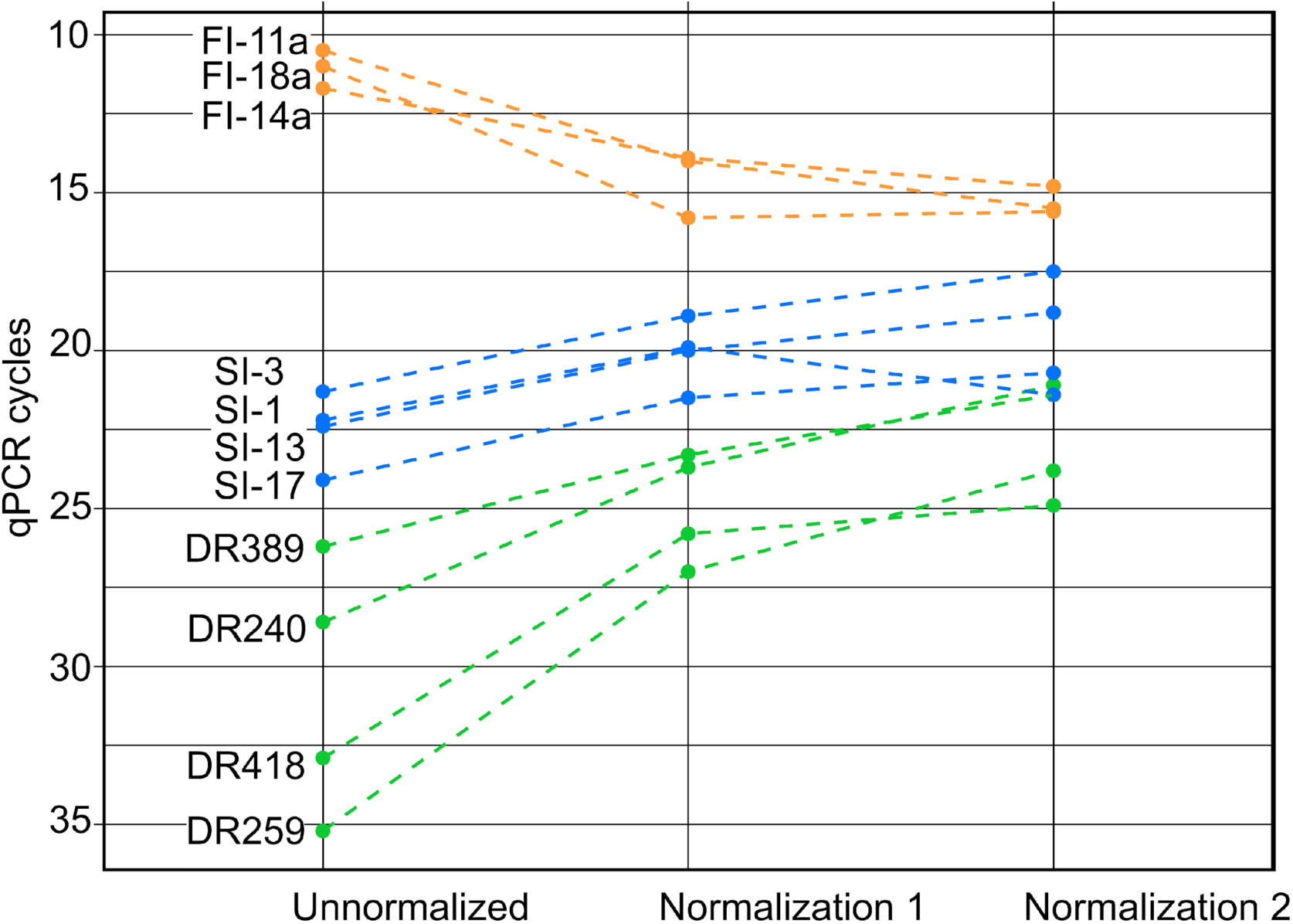
qPCR analysis of selected insertions and spike-in controls. Each dot indicates Ct values for each Alu flanking sequence in “unnormalized”, “normalization 1” and “normalization 2” libraries. Orange dots – fixed insertions (FI) present in each cell, blue dots – somatic insertions (SI) present in more than one cell, green dots (DR) – spike-in controls containing artificial sequences from *Danio rerio*. The difference in Ct between abundant fixed insertions and rare spike-in insertions changed from 25 cycles for “unnormalized” to 10 cycles for “normalization 2” libraries.

### Parameters of amplicon library normalization

More generally, the effect of normalization is described by the normalized entropy measure that evaluates distribution uniformity of sequencing reads per insertion (The normalized entropy equals one if each insertion is covered by an equal number of sequencing reads, and asymptotically approaches zero as the reads per insertion count becomes more biased). For the “unnormalized” library, the normalized entropy was estimated at 0.62. After the first and second rounds of normalization the entropy was increased up to 0.85 and 0.92 respectively. Thus we conclude that normalization makes the distribution of reads per insertions more even and increase the total number of different insertions detected, hence leading to the more efficient discovery of low represented insertions (see supplementary figure 1).

Renaturation of an amplicon during normalization is a complex process where many different types of molecules are hybridized to each other. For each group of molecules with the identical nucleotide sequence the speed of renaturation is mainly proportional to concentration although other factors including molecules length and GC content are also important. To evaluate the impact of these two factors on the normalization efficiency we plotted the number of sequencing reads corresponding to each Alu insertion from Ya5 (highly abundant before normalization) and Yb8 (rare before normalization) subfamilies versus the length of each fragment (Fig 3a). No relation between fragment length and normalization efficiency was observed. The impact of GC content on the normalization efficiency was more complex (Fig 3b). We observed a lower normalization rate for AT rich fragments during the first round of normalization. However, during the second round, the normalization rate for AT rich fragments was similar to their counterparts with higher GC content.

**Figure 3.**
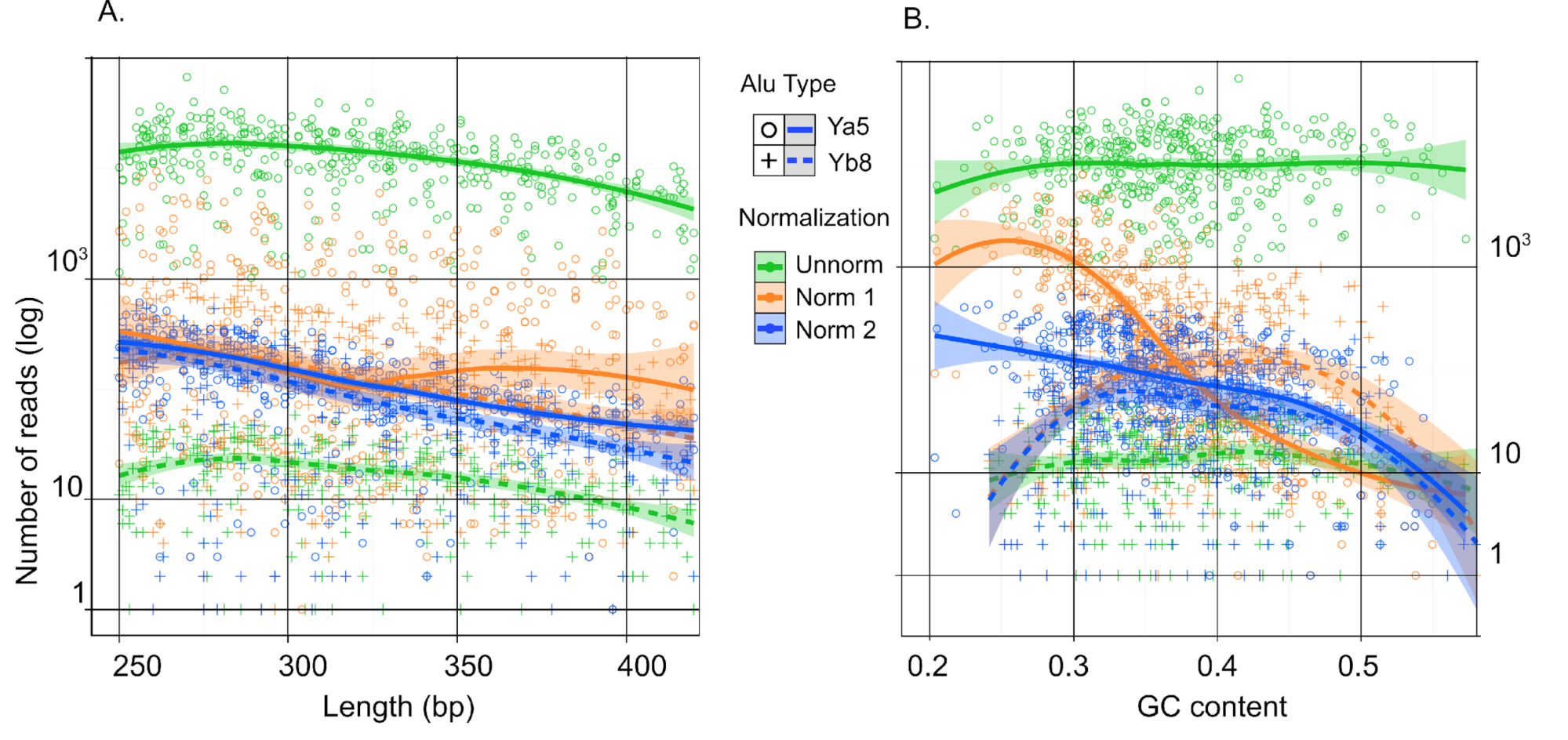
Effect of fragment length and GC content on normalization efficiency. The number of sequencing reads of rare (corresponding to AluYb8) and abundant (corresponding to AluYa5) flanks is plotted against fragment length (A) and GC content (B), respectively, in “unnormalized”, “normalization 1” and “normalization 2” datasets. Y axis – number of reads (logarithmic scale). X axis is length of fragments (A) or their GC content (B). Green circles and green crosses indicate Ya5 and Yb8 insertions in “unnormalized” library; orange circles and orange crosses indicate Ya5 and Yb8 insertions in “normalization 1” library; blue circles and blue crosses indicate Ya5 and Yb8 insertions in “normalization 2” library. Trendlines were fit to data using generalized additive models, shaded area indicate confidence interval (CI=0.95) for trendlines.

## Discussion

Somatic mosaicism resulting from new RE insertions was proposed to play a significant role in adult organism in particular contributing to individual neurons plasticity (8, 27). RE activity might also be involved in brain disorders including Rett syndrome (9) and schizophrenia (28). The most valid method to find new somatic RE insertions is their direct detection by high-throughput sequencing of genomic DNA. Although the capacity of modern sequencing platforms is rapidly increasing it is still expensive to study the distribution of somatic RE insertions (especially rare) in thousands of individual cells or many tissue samples. Even with the use of current protocols for enrichment in RE sequences only a minor fraction (up to 0.01% (13, 14)) of HTS reads is comprised by the somatic elements. In this study, we propose a tool that can significantly improve the capacity of most methods to identify rare somatic RE insertions. The entire process supposes two types of enrichment procedures: (1) selection of sequences flanking RE insertions of a particular subfamily by one of existing methods and (2) enrichment for sequences representing somatic insertions (normalization). The better results at the first enrichment stage are achievable using vectorett PCR or suppressive PCR techniques (13). As a result, more than 90% of the final amplicon is comprised by DNA fragments that flank RE insertions of the selected subgroup. During the second enrichment stage (employed in this study) highly abundant fragments are diminished in the amplicon, while rare sequences (including the fragments corresponding to somatic RE insertions) are enriched. Thus, two successive rounds of normalization led to more than 26-fold increase in the number of somatic REs flanks in a sequenced sample. The efficiency of this strategy is confirmed by both direct sequencing and qPCR of somatic insertions and spike-in controls.

Along with a more than 26-fold increase in the number of sequencing reads representing somatic REs, the number of identified somatic insertions increased by 7.9-fold (from 48 to 379) and the UMI number increased by 9.2-fold (from 51 to 468). The difference between the increments of the sequencing reads and somatic insertions might be explained by the limited number of somatic REs present in 50,000 cells. Therefore, the enrichment by normalization increases the number of reads, while the number of identified insertions starts reaching a plateau.

In this study we employed two successive rounds of normalization. The first normalization round resulted in a 10-fold increase in the number of sequencing reads corresponding to somatic insertions and 3.5-fold increase in the number of identified somatic insertions. After the second round of normalization there was an additional ~2.5-fold increase in both the number of reads and the number of somatic insertions. The difference in the efficiency of the first and second rounds of normalization probably reflects the principal limitation of the method of enrichment for low abundant fragments under selected conditions (renaturation time and DNA concentration).

UMIs are increasingly applied in the HTS-based methods to reduce the biasing effect of PCR and sequencing on quantitative information about particular sequences in the initial sample and to correct for PCR and sequencing errors (16, 17, 30). For instance, UMIs were used recently for the quantitative assessment of T cell repertoire diversity in course of aging (31). Although deep oversequencing is usually required for the accurate estimation of UMI based events (30), some unique quantitative traits could be obtained even with smaller sequencing depth. Here we ligate UMIs before introducing any quantitative bias by selective PCR or bridge amplification on the solid phase of the Illumina sequencing machine. Thus, the number of UMIs ligated to the fragments with identical sequences corresponds to the number of cells bearing this particular insertion.

Furthermore, the use of normalization in combination with UMI can help to distinguish various artifacts from the true somatic events. Formation of artifacts is a sporadic process which normally can’t be repeated even twice. As retroposition usually occurs in dividing cells, the new somatic insertion can be found at least in few cells and will have more than one UMI in contrast to chimeric molecules on condition of appropriate sequencing depth which can be reached by the use of normalization procedure. The similar technique based on signal strength differences between true insertions and artifacts was previously used in single cell based technique and now can be applied to artifact filtering in multiple cell sequencing approaches. In this study we found 39 potential somatic AluYa5 insertions (Table 2) which were characterised by more than one UMI per insertion. Therefore, these ones represent the most reliable pool of potentially somatic insertions detected in this study, however, the final validation can be reached only by identification of target site duplication (TSD) - the main characteristic signature of retroposition event. Simultaneous sequencing of both 5’ and 3’ RE flanks for TSD identification as well as the developed normalization based enrichment technique could significantly improve existing methods for the rare somatic RE insertions profiling.

## Conclusions

Somatic RE activity in humans and other mammals has been intensively studied over the last several years. Several studies reported a significant rate of insertional mutagenesis mediated by *de novo* integrations of REs not only in cancer, but also in normal human tissues including the brain. However, current enrichment protocols do not provide enough power for the detection of novel RE integrations and thus the sensitivity for somatic RE detection is usually enhanced by increasing the number of sequencing reads, which is cost consuming. The described approach can increase the efficiency of existing RE identification methods decreasing the number of sequencing reads required for the confident estimation of somatic REs abundance. Furthermore, the method allows to analyze much larger samples (tens of thousands cells) than usually studied nowadays (from 1 to hundreds of cells) with an almost comprehensive identification of very rare somatic RE insertions. The use of UMIs provides quantitative information on the distribution of REs. The direct estimation of the number of cells bearing each particular insertion can provide information on the period of RE retroposition activity in studied tissues, which could be linked to the stage of the disease progress or normal tissue development.

## Methods

### Nuclei isolation and DNA extraction

100 mg frozen tissue from postmortal human cortex (72 year old male individual) was used for nuclei isolation. All following manipulations were performed on ice. Tissue sample was homogenized in Dounce tissue grinder in 10 ml of nuclei extraction buffer (10 mM Hepes, 3 mM MgCl_2_, 5 mM CaCl_2_, 0.32 M sucrose, 0.2% Triton X-100). Homogenate was layered over equal volume of sucrose solution (0.64 M sucrose, 1×PBS, 0.2% Triton X-100) and centrifuged for 15 minutes at 1600g, +4°C. The sediment was resuspended in 1 ml 1×PBS and centrifuged for 10 minutes at 450g, +4°C. The obtained nuclei fraction was resuspended in 200 µl 1×PBS, stained by trypan blue and counted in cytometer. Genomic DNA extracted from approximately 50,000 nuclei was taken for downstream applications.

### AluYa5 flanking fragments library preparation

Genomic DNA was digested by incubation with AluI (Fermentas) endonuclease (10 U) for 12 hours. Fragmented DNA was purified by AmPure XP beads (Beckman Coulter) and ligated to suppressive adapters. The 10 µl ligation mixture contained 50 pmoles of each st19BH and st20BH adapters, 10 U of T4 DNA ligase in a T4 reaction buffer (both Promega) and digested genomic DNA. The reaction was carried out overnight at +4°C. Ligated fragments were incubated for 2 hours with 3 U of restriction enzyme AluI in 1× Y Tango buffer to decrease the number of chimeric molecules. Restriction products were purified using QIAquick PCR Purification Kit (Qiagen).

DNA amplification for library preparation was performed in two subsequent suppression PCR steps.

Each of 20 first step PCR reaction (25 µl) contained 1/20 of the total amount of ligation reaction, 0.4 µM AluYa5/AluYa8 specific primer (AY107), 0.16 µM Na21 primer, dNTP (0.125 µM each), 1 U of Tersus polymerase in 1× Tersus buffer 2 (both Evrogen). The amplification profile was as follows: 72°C for 4 min, followed by 12 cycles of 20 s at 94°C, 15 s at 65°C and 1 min at 72°C. PCR products were combined, purified with the QIAquick PCR. Each of two second step PCR reaction (25 µl) contained 1/160 of the first PCR products, 0.4 µM of each AY16-6 and st19okor primers, dNTP (0.5 µM each), 1 U of Tersus polymerase in 1× Tersus Plus buffer. The amplification profile was as follows: 20 s at 94°C, 15 s at 60°C, 1 min at 72°C, 9 cycles. PCR product was purified and loaded on agarose gel. Fragments ranging from 250 to 450 bp were cut and purified using QIAquick Gel Extraction kit (Qiagen).

### Spike-in controls preparation

Four different loci of zebrafish genome were selected for the preparation of artificial spike-in controls. Four different PCR reactions (25 µl) containing 20 ng of zebrafish genomic DNA, dNTP (0.125 µM each), 1 U of Tersus polymerase and 0.4 µM of each DR primers (see Supplementary table 1, primers for spike-in preparation) in 1× Tersus Plus buffer were performed. Forward primer contained the 16 nucleotides of AluYa5 at the 5ʹ end. The amplification profile was as follows: 20 s at 94°C, 15 s at 60°C, 1 min at 72°C, 9 cycles. Obtained PCR products were phosphorylated using T4 polynucleotide kinase (Promega) in the appropriate buffer. Phosphorylated PCR products were ligated to St19BH/St20BH adapter as described above. On the last step PCR reaction with ligated fragments and 0.4 µM of each AY16-6/St19okor primers was performed. PCR products were purified by Cleanup mini PCR Purification Kit (Evrogen) and their concentration was measured by Qubit. As a result four DNA fragments with the ends identical to those of the constructed AluYa5 flanking fragments library and having four different flanking sequences 240, 259, 389 and 418 bp long inside were obtained. 0.6×10^−9^ ng of DR259, 1×10^−9^ ng of DR418, 2.2×10^−9^ ng of DR240 and 3.6×10^−9^ ng of DR389 were added to 4.2 ng of AluYa5 flanking fragments library that corresponds to the insertions present in one (DR259 and DR418) or 5 (DR240 and DR389) out of 50,000 cells. AluYa5 flanking fragments library with added spike-in controls hereafter is called DNA mix 1.

### Normalization with Kamchatka Crab duplex-specific nuclease (DSN)

An aliquot (1/6 part) of the obtained DNA mix 1 were used for “unnormalized” control library preparation. Each of 5 PCR reaction tubes (25 µl) contained 1/30 of the DNA mix 1, 0.8 µM of each AY16-ind301 (contains sample barcode 301) and st19okor primers, 0.25 µM each of dNTP, 1 U of Encyclo polymerase in the 1× Encyclo reaction buffer (both Evrogen). The amplification profile was as follows: 9 cycles of 20 s at 94°C, 15 s at 60°C, 1 min at 72°C. PCR products were combined and purified using QIAquick PCR Purification Kit (Qiagen).

Same volume aliquot of DNA mix 1 was subjected to PCR as described above except for primers used for amplification (AY16-6 without sample barcode and st19okor, 13 cycles). 480 ng (3 µl) of the purified PCR product was mixed with 1 µl of 4× Hybridisation Buffer (200 mM HEPES pH 7.5, 2M NaCl). Reaction mixture was overlaid by mineral oil drop, denatured at 97°C for 3 min, chilled to 76°C with ramp 0.1°C/s and renatured at 76°C for 4 hours. After renaturation 5 µl of 2× DSN Master Buffer and 1 µl (1 U/µl) of DSN solution (both Evrogen), preheated at 76°C, were added to the reaction consequentially. Incubation was continued at 76°C for 15 min. 10 µl of 2× Stop Solution (Evrogen) was added to the reaction to inactivate DSN. The resulted normalization product was immediately purified using AMpure XP beads (Beckman Coulter, USA) and redissolved in 30 µl of water.

First aliquot (15 µl) was reamplified with AY16-ind302/st19okor primers and Encyclo polymerase for 9 cycles as described above resulting in “normalization 1” library. Second aliquot (15 µl) was reamplified with AY16-6/st19okor primers and used for second normalization as described above except of higher DNA concentration (1800ng in 3 µl). After the second normalization DNA was purified using AMpure XP beads and reamplified with AY16-ind304/st19okor primers and Encyclo polymerase for 9 cycles as described above resulting in “normalization 2” library.

### Sequencing and data analysis

Three libraries (“unnormalized”, “normalization 1” and “normalization 2”) each containing sample barcode were ligated to Illumina Truseq adapters using standard protocol and sequenced on HiSeq 2000 platform (paired end 2×100). Sequencing reads were analyzed using the pipeline described recently (13, 18) with the use of standard tools: Bowtie2 (19), Galaxy (20-22) and Python scripts for removal of artifacts (sequences of ligation chimeras, incorrect priming molecules and template switching molecules) from analyzed data set.

### Quantitative PCR analysis of selected AluYa5 insertions and spike-in controls

qPCR was performed to measure relative quantities of four fixed, four selected somatic and four artificial spike-in AluYa5 insertions. Each pair of primers was designed to align to unique gemomic region between 5ʹ end of the Alu element and nearest AluI restriction site. Each of 15 µl PCR reactions contained 2.5 ng of template DNA (“unnormalized”, “normalization 1” or “normalization 2” libraries), 0.17 µM of each direct and reverse primers (see Supplementary table 1, primers for qPCR) in 1× qPCR-HS SYBR mix (Evrogen). Three technical replicates for each PCR reaction were performed. The changes in relative quantities were evaluated using delta-delta Ct method.

### Data processing and statistical analysis

To evaluate the statistical significance of discovering new somatic insertions we applied Pearson’s Chi-squared test. The P values were calculated using the chisq.test function from R (23). The normalized entropy measure on a distribution of reads per insertion *D* for a sample was calculated using the following formula:

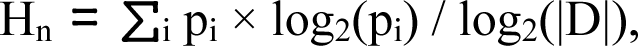

where Hn is normalized entropy, p_i_ is a proportion of reads in the i-th insertion to the overall number of reads, |D| is a size of the distribution (total number of identified insertions). To correct sequencing errors in UMIs corresponding to each somatic Alu insertion we built a graph where UMI sequences were vertices and hamming distances between them were edges. Each strongly connected component in the graph with one “parental” UMI was deleted. Number of remaining vertices was considered as a corrected number of UMIs in the input set for each particular somatic RE insertion.

### Filtering of chimeric sequences

Both obvious and hidden chimera sequences was removed from HTS data sets after genome mapping. Sequencing reads discordantly mapped on the genome was considered as obvious chimeras. Reads mapped properly but on loci containing restriction site or sequence similar to known AluYa5 flanks in proximity of putative integration point was considered as hidden chimeras produced by ligation and PCR template switching respectively. Sequences produced during incorrect priming during PCR was also removed from the data set.

### Accession numbers

The raw Illumina sequencing data have been deposited in the NCBI Sequence Read Archive (SRA) with accession number SRX1113412.

## List of abbreviations

RE: retroelement
UMI: unique molecular identifier
qPCR: quantitative polymerase chain reaction
HTS: high-throughput sequencing

## Declarations

The study was approved by the local ethics committee and conducted in accordance with the Declaration of Helsinki.

This work was supported by Russian Foundation for Basic Research grants [RFBR 16-04-00779 to IZM, RFBR 16-34-01100 and 17-04-01280 to AYK], Russian Science Foundation grants [RSF 17-75-10113 to AYK and RSF 18-14-00244 to IZM] and Russian President’s Fellowship [SP-4059.2016.4 to AYK].

---

**Supplementary figure 1.** Distribution of reads per insertion in “unnormalized”, “normalization 1” and “normalization 2” libraries. X axis – reads per insertion (log scale), Y axis – number of insertions covered by particular number of reads (square root of proportion)

